## Supplementary Information for "An Hfq-dependent post-transcriptional mechanism fine tunes RecB expression in *Escherichia coli*"

#### Supplementary Notes

#### A stochastic two-stage model of RecB expression

To model RecB expression, we applied a well-established stochastic mRNA-protein model of gene expression and inferred its parameters by fitting to the experimental data. Here, we briefly introduce the model, discuss its underlying assumptions and provide the analytical expressions used in the fitting procedure.

**Model description**

A stochastic two-stage model of gene expression has been widely utilized to study fluctuations underlying protein production in prokaryotic and eukaryotic cells [1, 2, 3, 4, 5, 6]. Generally, the model is based on four biological processes: transcription of a gene by an RNA polymerase, translation of an mRNA by a ribosome, mRNA degradation and protein decay. As elsewhere, we assume a cell to be a well-stirred system of biochemical molecules, the interaction between which can be described with a common approach of chemical reactions [7]. We focus on two species, mRNAs and proteins, and denote molecule numbers in a cell as $m$ and $p$, respectively.

Considering gene expression in bacterial cells allows several further assumptions to be made. First, transcription of bacterial genes was demonstrated to occur in bursts [8, 9] and quite a few transcriptional models describing gene bursting have been proposed in the literature [3, 10, 11, 12]. Thus, mRNA production is described as a zeroth-order chemical reaction where a gene is transcribed in mRNA bursts with a constant rate $k_m$ and the number of mRNAs per each burst $n$ is sampled from a geometric distribution with a mean size $b$ [13, 14, 15]. Secondly, based on a broadly accepted view of the exponential RNA degradation in bacteria [16, 17, 18], mRNA decay is modelled as a first order reaction with a constant rate $\gamma_m$. Furthermore, proteins in bacteria were shown to be diluted exclusively because of cell growth and division [19, 20]. This process can be implicitly modelled by a first-order chemical reaction with a constant rate equal to growth rate $\gamma_p$. Finally, we assume that a protein is produced from an mRNA with a constant rate $k_p$. Thus, the entire reaction network for the two-stage model with burst transcription is summarized in the following schematic:

$$\begin{aligned} &\varnothing \xrightarrow{k_m} n\,m \qquad && m \xrightarrow{k_p} m + p \\ &m \xrightarrow{\gamma_m} \varnothing \qquad && p \xrightarrow{\gamma_p} \varnothing \end{aligned} \tag{A1}$$

The dynamics of the mRNA and protein average levels (denoted as $\langle m \rangle$ and $\langle p \rangle$) is described by a set of the following rate equations:

$$\frac{d\langle m \rangle}{dt} = b\,k_m - \gamma_m \langle m \rangle \tag{A2}$$

$$\frac{d\langle p \rangle}{dt} = k_p \langle m \rangle - \gamma_p \langle p \rangle \tag{A3}$$

**Steady-state mRNA and protein mean and variance**

The stationary solutions for mean mRNA and protein numbers can be found by imposing the steady-state conditions ($d\langle m \rangle/dt = 0$ and $d\langle p \rangle/dt = 0$) to the rate equations (Eq. (A2) & Eq. (A3)):

$$\langle m \rangle = b\frac{k_m}{\gamma_m} \qquad \langle p \rangle = b\frac{k_m k_p}{\gamma_m \gamma_p} \tag{A4}$$

As a result of the linearity of the system, the mRNA and protein variances can be found exactly by van Kampen's Ω-expansion [7, 21, 22].

$$\sigma_m^2 = (1+b)\langle m \rangle \qquad \sigma_p^2 = \langle p \rangle + \frac{k_p^2(1+b)\langle m \rangle}{\gamma_p(\gamma_m+\gamma_p)} \tag{A5}$$

Finally, the mRNA and protein coefficients of variation (squared), defined as variance over squared mean, are given as follows [1, 2, 3, 23]:

$$CV_m^2 = \frac{\sigma_m^2}{\langle m \rangle^2} = \frac{1+b}{\langle m \rangle} \tag{A6}$$

$$CV_p^2 = \frac{\sigma_p^2}{\langle p \rangle^2} = \frac{1}{\langle p \rangle} + \frac{1+b}{\langle m \rangle} \frac{\gamma_p}{\gamma_m + \gamma_p} \tag{A7}$$

The analytical probability distribution of protein numbers under assumption of a long-lived protein relative to short-lived mRNA was derived here [1].

#### Deterministic model of RecB expression under DNA damage conditions

In this section, we analytically describe the dynamics of RecB mRNA and protein levels after a perturbation made by DNA damage with a sub-lethal dose of ciprofloxacin. Taking into account different time-scales of the response in mRNA and protein levels, we address the following question: how long does the DNA damage need to be applied for to see changes (if any) in both species? We focus on the average mRNA and protein concentrations instead of molecule numbers because of cell elongation upon ciprofloxacin treatment (Figure 3B & 3C) and modify a two-stage model of RecB expression by including deterministic time-dependency of cell volume, $V(t)$.

As discussed above, we consider the same set of reactions for two species of the system: mRNAs and proteins. While the processes of translation, active mRNA and protein degradation are described by first-order reactions with constant rates: $k_p$, $\gamma_m$ and $\gamma_{p(act)}$, respectively, mRNA transcription is given by a zeroth-order reaction. According to the law of mass action, we shall scale a zeroth-order reaction rate by the volume of the system, as $k_m V(t)$ [7, 24]. We define the number of mRNA and protein molecules at time $t$ in a cell of volume $V(t)$ as $m(t)$ and $p(t)$, respectively. Then, the evolution of the average numbers of mRNA and protein molecules in time is described by the following rate equations:

$$\frac{d\langle m(t)\rangle}{dt} = k_m V(t) - \gamma_m \langle m(t)\rangle \tag{A8}$$

$$\frac{d\langle p(t)\rangle}{dt} = k_p \langle m(t)\rangle - \gamma_{p(act)} \langle p(t)\rangle \tag{A9}$$

We further define *recB* mRNA and RecB protein concentration as $c_m(t) = m(t)/V(t)$ and $c_p(t) = p(t)/V(t)$ and differentiate with respect to $t$:

$$\frac{dc_m(t)}{dt} = \frac{1}{V(t)} \frac{dm(t)}{dt} - \frac{c_m(t)}{V(t)} \frac{dV(t)}{dt} \tag{A10}$$

$$\frac{dc_p(t)}{dt} = \frac{1}{V(t)} \frac{dp(t)}{dt} - \frac{c_p(t)}{V(t)} \frac{dV(t)}{dt} \tag{A11}$$

In order to obtain time-dependent solutions for mRNA and protein concentrations, we divide the rate equations (Eq. (A8) & Eq. (A9)) by cell volume $V(t)$ and replace $dm(t)/dt$ and $dp(t)/dt$ with the expressions from the equations (Eq. (A10) & Eq. (A11)). Based on the experimental evidence [25, 26], we assume the exponential dependence of cell volume from time, $V(t) = V_0 e^{\gamma t}$, where $\gamma$ is a growth rate. Thus, we obtain the following ordinary differential equations for mRNA and protein concentrations:

$$\frac{d\langle c_m(t)\rangle}{dt} = k_m - (\gamma_m + \gamma) \langle c_m(t)\rangle \tag{A12}$$

$$\frac{d\langle c_p(t)\rangle}{dt} = k_p \langle c_m(t)\rangle - (\gamma_{p(act)} + \gamma) \langle c_p(t)\rangle \tag{A13}$$

By equating the left-hand sides of the equations to zero, we find the steady-state mRNA and protein concentrations, $c_m^{ss}$ and $c_p^{ss}$:

$$c_m^{ss} = \frac{k_m}{\gamma_m + \gamma} \qquad c_p^{ss} = \frac{k_m}{(\gamma_m + \gamma)} \frac{k_p}{(\gamma_{p(act)} + \gamma)} \tag{A14}$$

Next, by assuming the initial conditions for mRNA and protein concentration to be $c_{m(t=0)}$ and $c_{p(t=0)}$, we obtain the time-dependent solutions for mRNA and protein concentrations:

$$\langle c_m(t)\rangle = \left(c_{m0} - \frac{k_m}{\gamma_m+\gamma}\right) e^{-(\gamma_m+\gamma)t} + \frac{k_m}{\gamma_m+\gamma} \tag{A15}$$

$$\langle c_p(t)\rangle = \left(c_{p0} + \frac{c_{m0}k_p}{\gamma_m - \gamma_{p(act)}} - \frac{k_p k_m}{(\gamma_m - \gamma_{p(act)})(\gamma_{p(act)}+\gamma)}\right) e^{-(\gamma_{p(act)}+\gamma)t} + \tag{A16}$$

$$+\left(-\frac{c_{m0}k_p}{\gamma_m - \gamma_{p(act)}} + \frac{k_p k_m}{(\gamma_m - \gamma_{p(act)})(\gamma_m+\gamma)}\right) e^{-(\gamma_m+\gamma)t} + \frac{k_p k_m}{(\gamma_{p(act)}+\gamma)(\gamma_m+\gamma)}$$

The last expression can be simplified by taking into account that the RecB protein does not have an active degradation mechanism (Figure 2D). Thus, after imposing the condition of $\gamma_{p(act)} = 0$, we obtain a time-dependent solution for RecB protein concentration as follows:

$$\langle c_p(t)\rangle = \left(c_{p0} + \frac{c_{m0}k_p}{\gamma_m} - \frac{k_p k_m}{\gamma_m \gamma}\right) e^{-\gamma t} + \left(-\frac{c_{m0}k_p}{\gamma_m} + \frac{k_p k_m}{\gamma_m(\gamma_m+\gamma)}\right) e^{-(\gamma_m+\gamma)t} + \frac{k_p k_m}{\gamma(\gamma_m+\gamma)} \tag{A17}$$

From the expressions Eq. (A15) & Eq. (A17) we can now estimate characteristic time-scales needed for mRNA and protein levels to change to the new conditions. It is worth noting that although cells are elongated upon ciprofloxacin treatment, they do not slow down the growth (Figure S5). Thus, the analysis of the obtained equations shows that mRNA solution has one mode, $e^{-(\gamma_m+\gamma)t}$, and hence a characteristic time-scale can be estimated as $1/(\gamma_m+\gamma) \sim 1.6\ min$ (the results of growth rate measurements, $\gamma = \gamma_{cipro^+} = 0.0165\ min^{-1}$, are shown in Figure S5c). In contrast, the expression for protein concentration is given by two time-scales: $e^{-\gamma t}$ and $e^{-(\gamma_m+\gamma)t}$, with the slowest one $1/\gamma \sim 61\ min$. This means that the perturbation needs to be applied for at least $\sim 1$ hour in order to be able to detect changes in protein concentrations. In our experiments, DSBs were induced for two hours to guarantee fulfillment of the time-scale requirement (Figure 3).

### Supplementary Figures

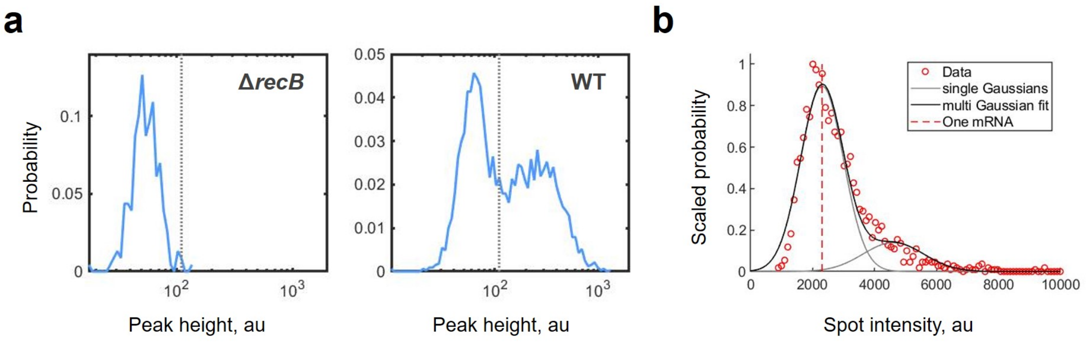


**Figure S1: smFISH image analysis performed by Spätzcell. a:** Peak height intensity profiles for spots detected in the wild-type and Δ*recB* samples. The dashed line indicates the intensity threshold (99.9 %) that separates a specific signal from the background. **b:** Integrated spot intensity histogram for foci detected in the wild-type sample. The data, binned and averaged in groups, are shown in red circles. The data were fitted by the sum of two Gaussian distributions corresponding to one and two mRNA molecule(s) per focus. The grey lines correspond to single Gaussian distributions, while the black solid curve is the sum of these two Gaussians. The red dashed line indicates the intensity equivalent corresponding to the total integrated intensity of FISH probes (in average) bound to a single mRNA (One mRNA).

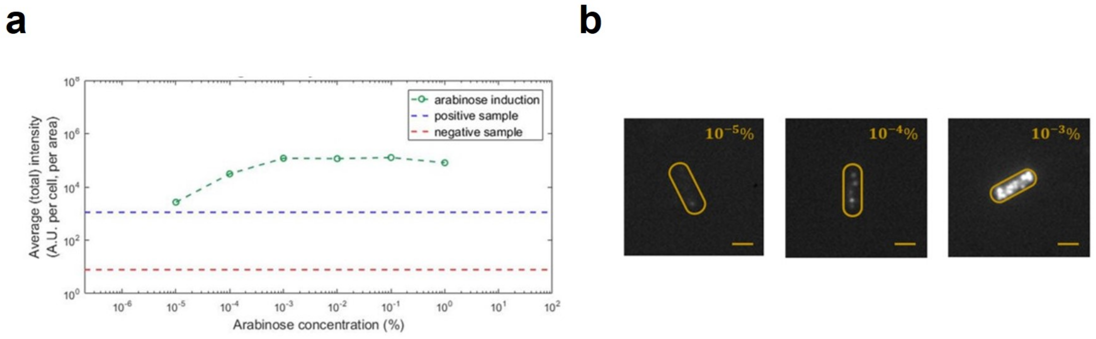


**Figure S2: Sensitivity of the smFISH protocol allows for quantification of low-abundant *recB* mRNAs. a:** Fluorescence signal, detected in the samples with *recB* over-expression from an arabinose-inducible plasmid (pIK02), is presented as a function of arabinose concentration (green-dashed line). The intensity of the fluorescence signal in the wild-type and Δ*recB* strains is indicated with blue-dashed and red-dashed lines, respectively. More than 350 cells for each induction condition were taken for the analysis. **b:** Representative fluorescence images of the samples where *recB* expression was induced with $10^{-3}$%, $10^{-4}$% or $10^{-5}$% of arabinose. All fluorescence images are shown in the same intensity range. The yellow line segment represents 1 $\mu m$.

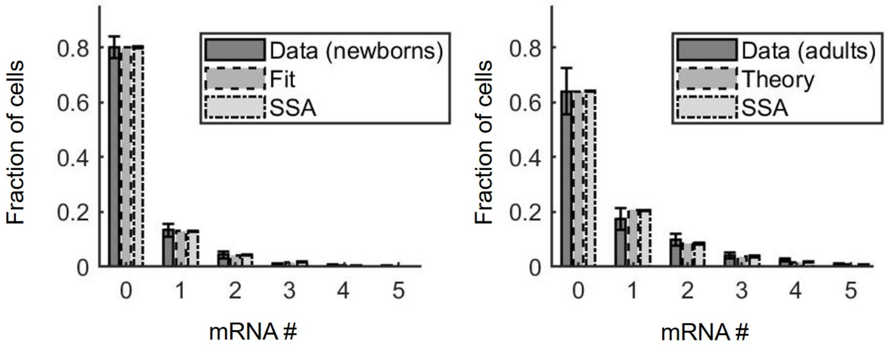


**Figure S3: Independent transcription from *recB* gene copies.** *recB* mRNA distributions for *newborns* (**left**) and *adults* (**right**). The experimental data from Figure 1C was conditioned on cell size (< 3.0 $\mu m^2$ for *newborns*) and (> 3.5 $\mu m^2$ for *adults*). Total number of cells: 2,180 (*newborns*) and 6,413 (*adults*). The *recB* mRNA statistics for *newborns* is well described by a negative binomial distribution, $NB(r, p)$ (Figure 1F). The sum of two independent variables, each of which is distributed as $NB(r, p)$, is then described by a negative binomial distribution $NB(2r, p)$. Thus, the *adults* data was compared to the predicted distribution, $NB(2r, p)$, based on (i) the assumption of independent transcription from *recB* gene copies and (ii) the fitting parameters inferred on the *newborns*. The results were verified with Gillespie's simulations (SSA). The Kullback–Leibler divergence between the experimental and simulated distributions for *newborns* and *adults* is $D_{KL} = 0.003$ and $D_{KL} = 0.007$, respectively.

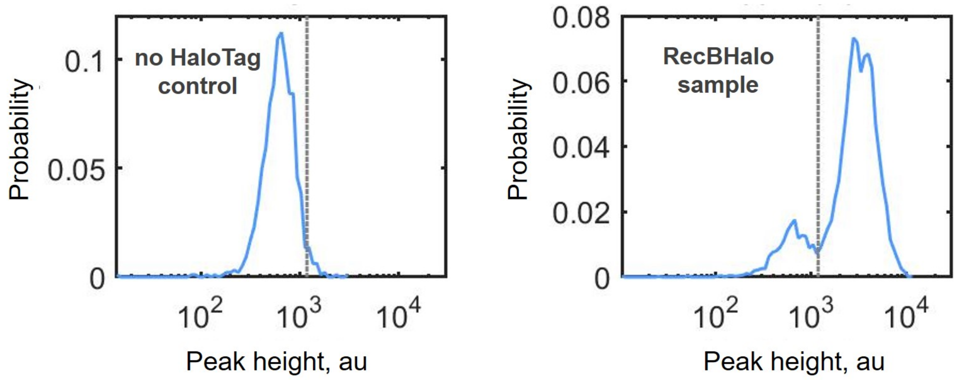


**Figure S4: Image analysis of Halo-labelling experiments performed with the modified version of Spätzcell.** Peak height intensity profiles for spots detected in the strain with RecB-HaloTag fusion (RecBHalo sample) and its parental strain (no HaloTag control). The dashed line indicates a chosen intensity threshold in order to separate a specific signal from the background.

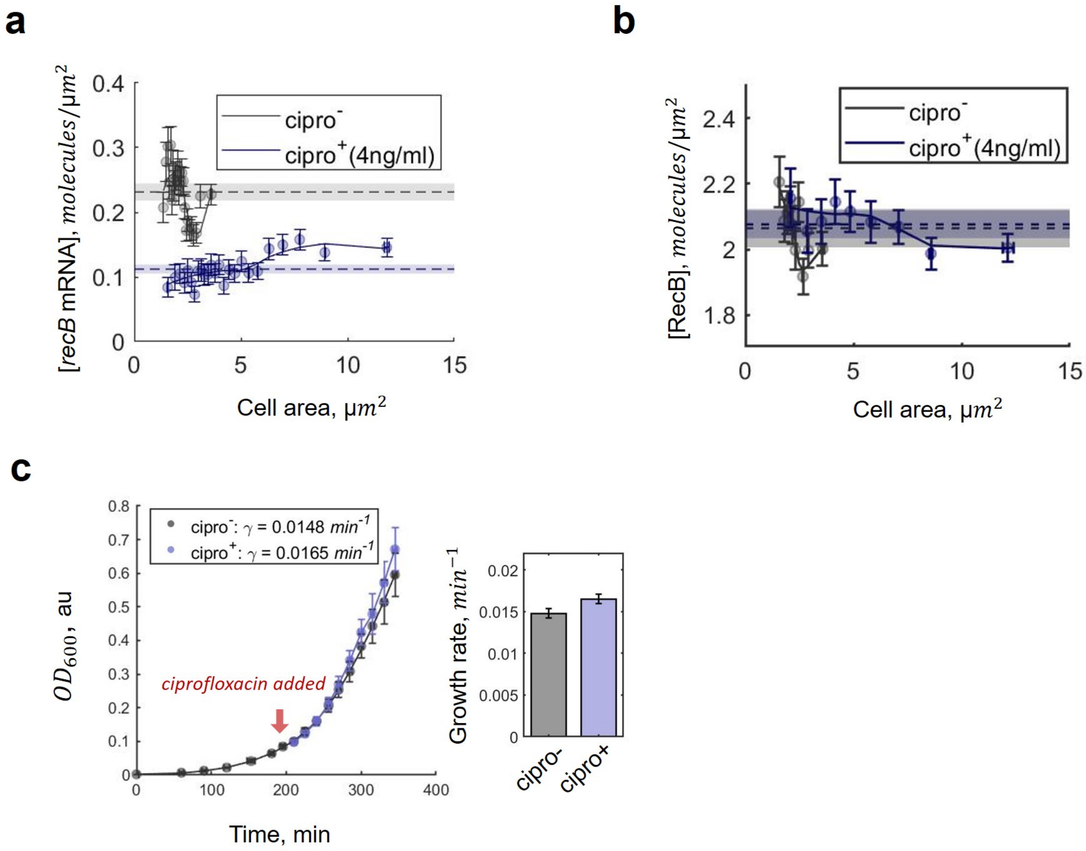


**Figure S5: Analysis of *recB* mRNA and RecB protein concentrations and growth rate measurements under DNA damage**. **a:** *recB* mRNA concentration for perturbed (blue) and unperturbed (grey) samples represented as a function of cell area. The data from Figure 3D were binned by cell size and averaged in each group (mean ± s.e.m.). The solid lines connect the averages while the dashed lines and shaded areas show mean ± s.e.m. calculated across all cells in each sample. **b:** RecB protein concentration for perturbed (blue) and unperturbed (grey) samples shown as a function of cell area. The data from Figure 3E were binned by cell size and averaged in each group (mean ± s.e.m.). The solid lines connect the averages while the dashed lines and shaded areas show mean ± s.e.m. calculated across all cells in each sample. **c:** Growth curves obtained with optical density measurements ($OD_{600}$). Cell cultures were grown with 4 ng/ml (blue) or without (grey) ciprofloxacin. The red arrow indicates the time when ciprofloxacin was added to the $cipro^+$ sample. The exponential phase of growth ($OD_{600} \sim$ [0.08-0.4]) was used for growth rate estimation. The growth rates, averaged across three replicated experiments, are shown in the legend. Error bars on the bar chart represent 95 % confidence intervals.

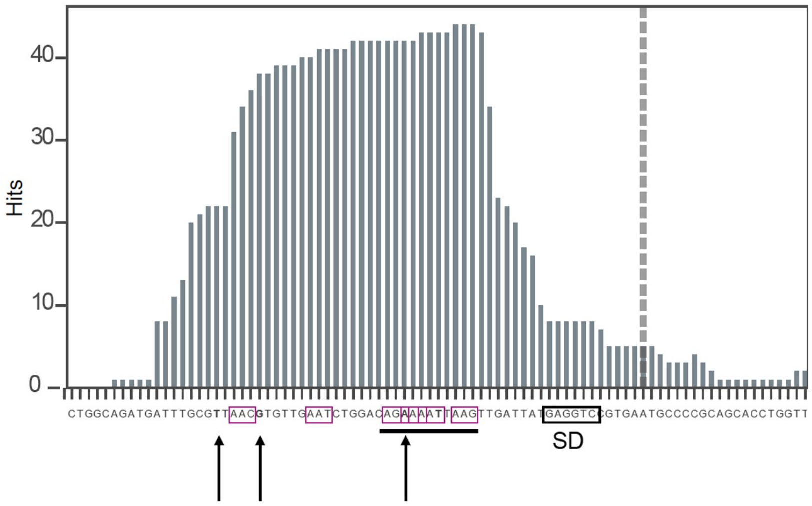


**Figure S6: The main Hfq binding site within *ptrA* gene.** The Hfq binding peak located within *ptrA-recB-recD* mRNA, identified in the Hfq-CRAC experiment [27], presented in raw counts (Hits). The vertical dashed line and black box indicate a translation initiation site (ATG) and Shine-Dalgarno sequence (SD), respectively. The black vertical arrows show the nucleotides where direct Hfq cross-linking was detected in the CRAC data. Hfq-binding motifs, A-R(A/G)-N(any nucleotide) [28, 29], are highlighted in the red boxes while the cluster of Hfq binding motifs is underlined.

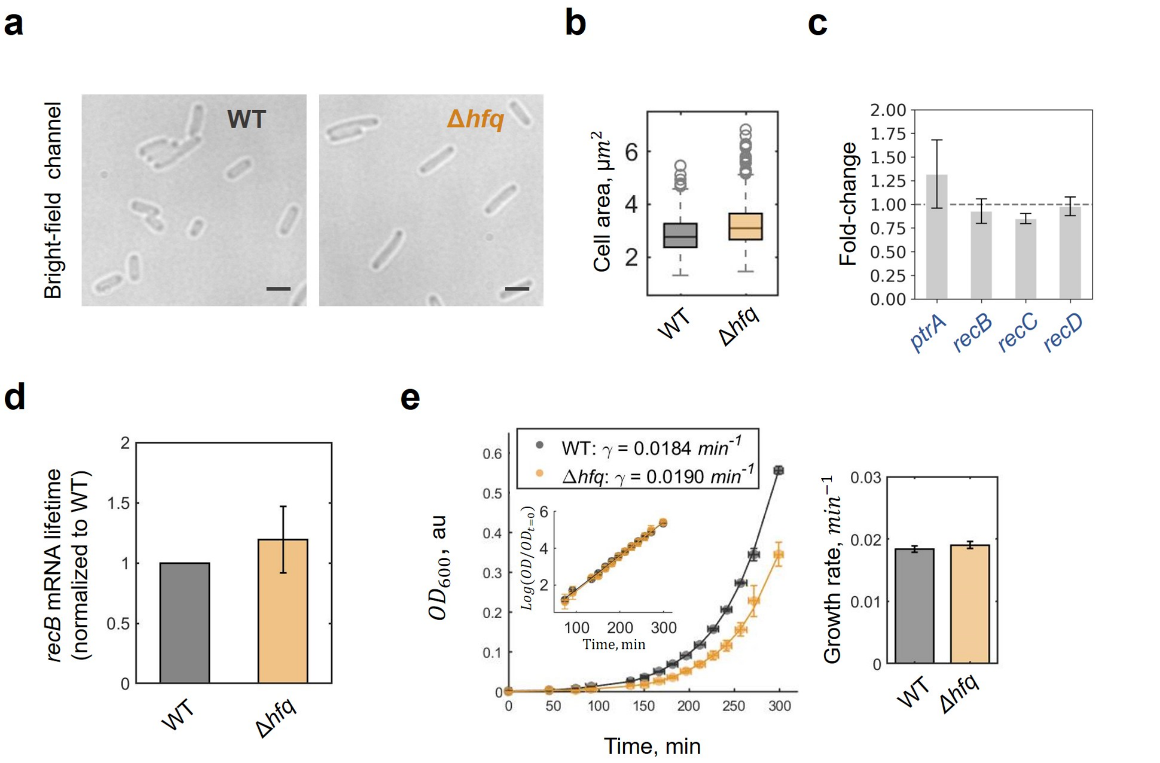


**Figure S7: Cell size analysis, RT-qPCR quantification and growth rate measurements in the Δ*hfq* mutant.** **a:** Examples of bright-field images for wild-type and Δ*hfq* strains. Scale bars represent 2 $\mu m$. **b:** Box plots represent cell area distributions for wild-type and Δ*hfq* samples. **c:** *ptrA*, *recB*, *recC* and *recD* transcripts quantified by RT-qPCR in the Δ*hfq* mutant and normalized to the corresponding expression levels in wild-type cells. The data represent averages and standard deviations across three replicates for each gene. *rrfD* was used as a reference gene. **d:** *recB* mRNA lifetime measured in the wild-type and Δ*hfq* mutant in a time-course experiment where transcription initiation was inhibited with rifampicin. mRNA abundance was quantified with RT-qPCR at t = 0, 1, 2 and 4 min after rifampicin treatment. *recB* mRNA lifetime was calculated from an exponential fit and normalized to the lifetime in the wild-type. The bar chart represents results of three replicated experiments; the error bar shows standard deviation. Significance was calculated with two-sample *t*-test (*P* value 0.35 (ns)). **e:** Growth curves obtained by optical density measurements ($OD_{600}$) in Δ*hfq* (orange) and wild type (grey). The strains were grown in the medium supplemented with glucose and amino acids. The exponential phase of growth ($OD_{600} \sim$ [0.08-0.4]), shown in the logarithmic scale in the insert, was used for growth rate calculation. The growth rates, averaged across three replicated experiments, are shown in the legend. Error bars on the bar chart represent 95 % confidence intervals.

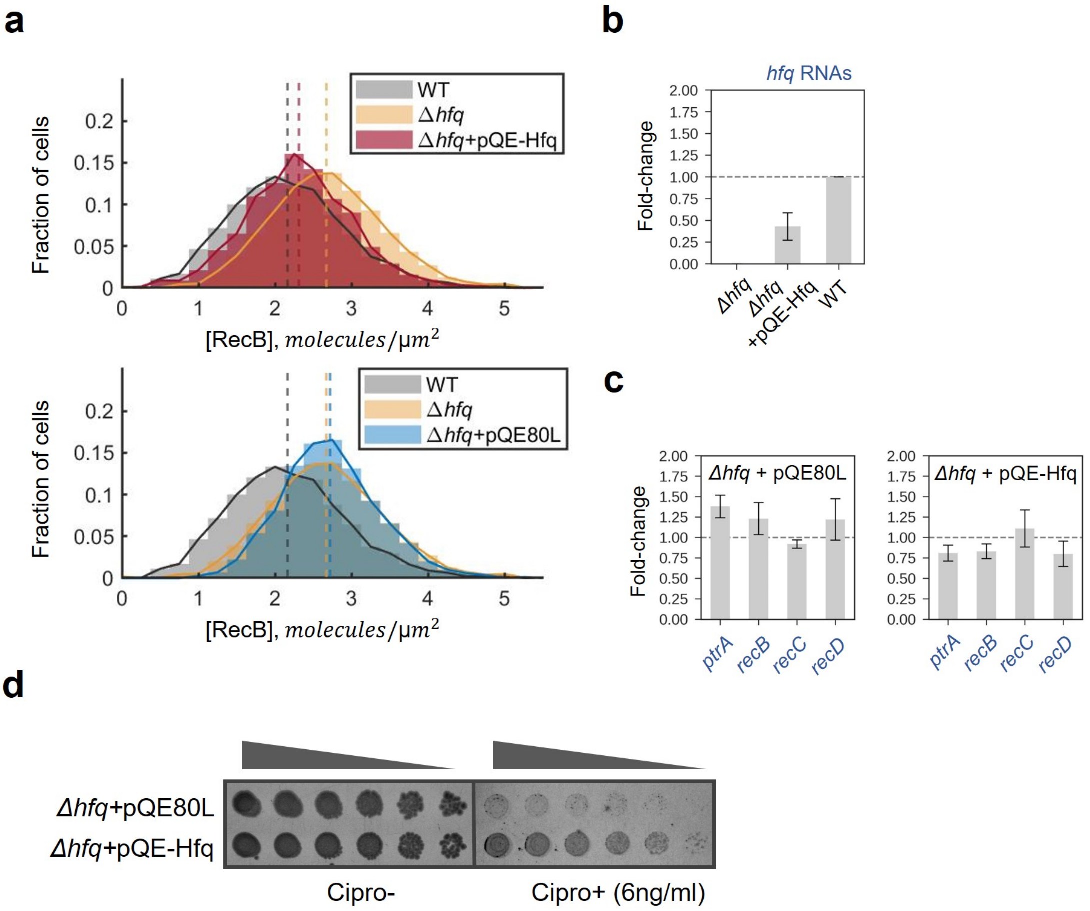


**Figure S8: Controls for Hfq complementation experiment.** **a:** RecB protein concentration distributions quantified in the Δ*hfq* mutant carrying pQE80L (blue) or pQE-Hfq (red) plasmid. The upper panel is the same as in Figure 5B and presented here to facilitate visual comparison with the lower panel. The histograms represent the average across two replicated experiments per each condition. The RecB concentration histograms for wild type and Δ*hfq* mutant are plotted as references in grey and orange, respectively. The dashed lines represent the mean RecB concentration in each condition. Significance was evaluated with two-sample *t*-test (*P* values: 0.045(*) for Δ*hfq* and Δ*hfq*+pQE-Hfq; 0.71(ns) for Δ*hfq* and Δ*hfq*+pQE80L). **b:** *hfq* RNA levels quantified by RT-qPCR in Δ*hfq* cells carrying pQE-Hfq plasmid and normalized to the *hfq* RNA level in wild-type cells. **c:** *ptrA*, *recB*, *recC* and *recD* transcripts quantified by RT-qPCR in Δ*hfq* carrying pQE80L (left) or pQE-Hfq (right) plasmid and normalized to the corresponding expression levels in Δ*hfq* and Δ*hfq*+pQE80L, respectively. **d:** A 5-fold serial dilution assay of Δ*hfq* strain carrying pQE80L or pQE-Hfq plasmid. Cells were plated onto LB plates without or with 6 ng/ml of ciprofloxacin.

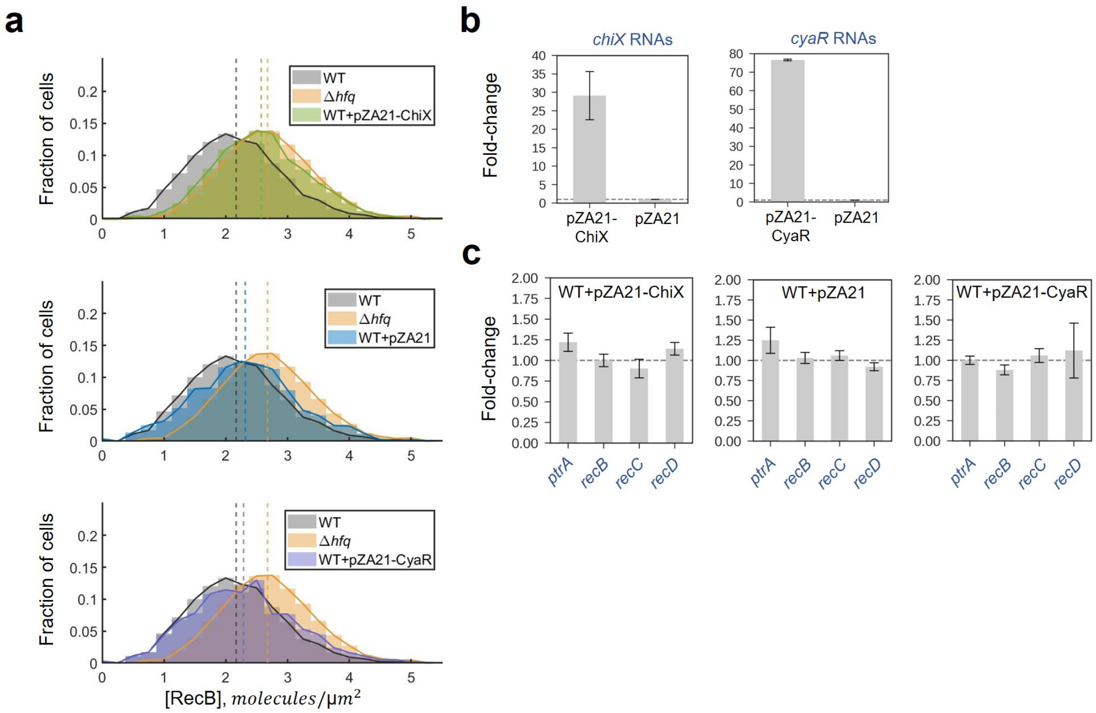


**Figure S9: Controls for ChiX over-expression experiment. a:** RecB protein concentration distributions quantified in wild-type cells carrying pZA21-ChiX (green), pZA21 (blue) or pZA21-CyaR (purple) plasmid. The upper panel is the same as in Figure 5C and presented here to facilitate visual comparison with the lower panels. The histograms represent the average across two replicated experiments per each condition. The RecB concentration histograms for wild type and the Δ*hfq* mutant are plotted as references in grey and orange, respectively. The dashed lines represent the mean RecB concentration in each condition. Significance was evaluated with two-sample *t*-test (*P* values: 0.021(*) for WT and WT+pZA21-ChiX; 0.42(ns) for WT and WT+pZA21; 0.37(ns) for WT and WT+pZA21-CyaR). **b:** *chiX* and *cyaR* RNA levels quantified by RT-qPCR in wild-type cells carrying pZA21-ChiX (left) or pZA21-CyaR (right) over-expression plasmid. The results were normalized to the corresponding expression in the cells carrying the backbone plasmid (pZA21). **c:** *ptrA*, *recB*, *recC* and *recD* transcripts quantified by RT-qPCR in wild-type cells carrying pZA21-ChiX (left), pZA21 (middle) or pZA21-CyaR (right) plasmid. RNA levels were normalized to the corresponding expression levels in wild type.

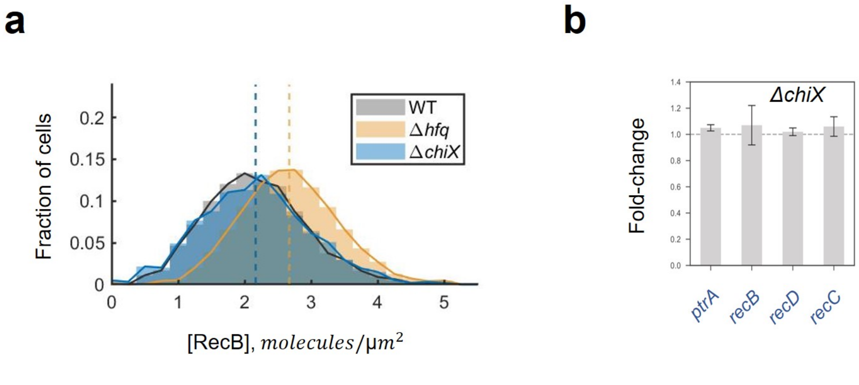


**Figure S10: RecB protein and mRNA quantification in Δ*chiX*. a:** RecB protein concentration distribution quantified in Δ*chiX* cells. The histogram represents the average across two replicated experiments. The RecB concentration histograms for wild type and the Δ*hfq* mutant are plotted as references in grey and orange, respectively. The dashed lines represent the mean RecB concentration in each condition. Significance was evaluated with two-sample *t*-test: *P* value for WT and Δ*chiX* is 0.96 (ns). **b:** *ptrA*, *recB*, *recD* and *recC* transcripts quantified by RT-qPCR in *chiX* mutants and normalized to the corresponding expression levels in the wild-type sample.

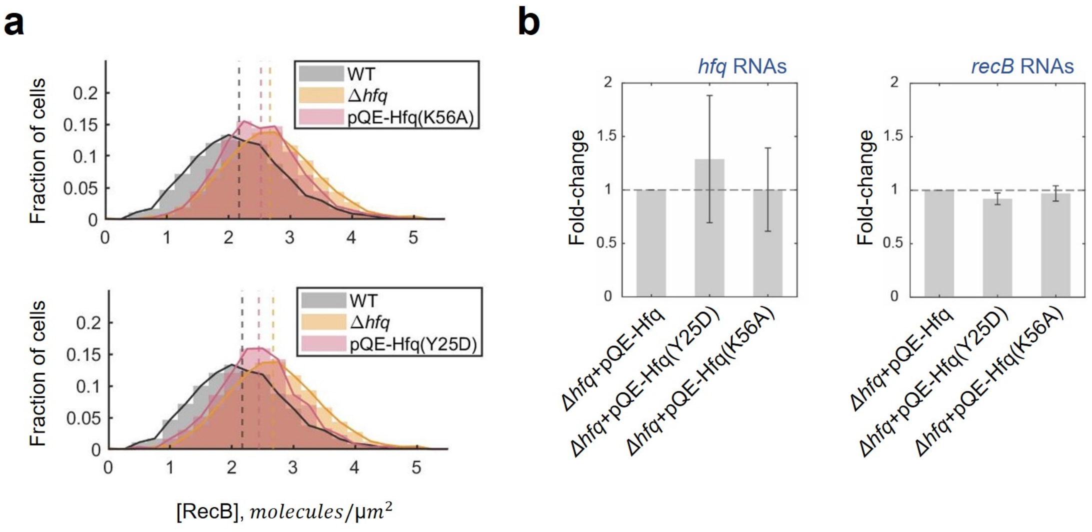


**Figure S11: RecB quantification upon expression of Hfq with point mutations in proximal or distal binding faces. a:** RecB protein concentration distributions quantified in *hfq* mutants carrying a pQE80L-derivative plasmid with Hfq protein mutated either in proximal (K56A) or distal (Y25D) binding faces [30]. The histograms represent the average across two replicated experiments for each mutation. The RecB concentration histograms for wild type and *hfq* mutants are plotted as references in grey and orange, respectively. The dashed lines represent the mean RecB concentration in each condition. Significance was evaluated with two-sample *t*-test: *P* values: 0.30(ns) for Δ*hfq* and Δ*hfq*+pQE-Hfq(K56A); 0.07(ns) for Δ*hfq* and Δ*hfq*+pQE-Hfq(Y25D). **b:** *hfq* and *recB* RNA levels quantified by RT-qPCR in Δ*hfq* cells carrying a pQE80L-derivative plasmid with a mutated Hfq protein. The results were normalized to the corresponding expression in Δ*hfq* cells carrying the backbone plasmid, pQE-Hfq.

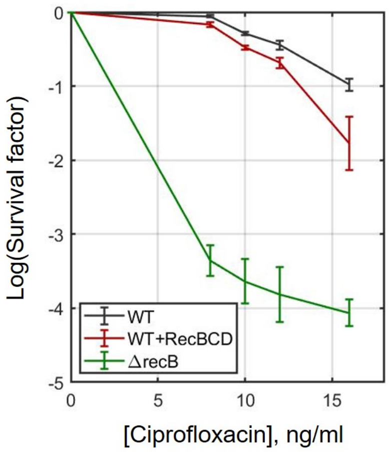


**Figure S12: Toxicity of RecBCD overexpression upon DSB induction.** Viability assays performed for the wild type, wild type carrying RecBCD over-expression plasmid, pDWS2, and Δ*recB*. Cells were plated onto LB plates supplemented with ampicillin and either without or with 8/10/12/16 ng/ml of ciproflaxacin. The average survival factors were calculated for at least three replicated experiments while the error bars indicate standard estimation of the mean.

#### Supplementary Tables

**Table S1:** *E. coli* strains used in the study.

| **Strain** | **Genotype** | **Source** |
|---|---|---|
| MG1655 | $F^- \lambda^-$ *ilvG*$^-$ *rfb-50 rph-1* | Lab stock |
| BW27783 | $F^- \lambda^-$ Δ*(araD-araB)567* Δ*lacZ4787*::rrnB-3 Δ*(araH-araF)570(::FRT)* Δ*araEp-532::FRT* $\phi P_{cp8}$*araE535 rph-1* Δ*(rhaD-rhaB)568 hsdR514* | Lab stock |
| MEK65 | MG1655 *recB165*::*halotag* | [31] |
| MEK1326 | MG1655 Δ*recB* | [31] |
| MEK1329 | BW27783 Δ*recB* | This work |
| MEK1902 | MG1655 Δ*hfq* | This work |
| MEK1457 | MG1655 *recB165*::*halotag* Δ*hfq* | This work |
| MEK1888 | MG1655 Δ*chiX* | This work |
| MEK1449 | MG1655 *recB165*::*halotag* Δ*chiX* | This work |
| MEK1938 | MG1655 *recB165*::*halotag* 5′UTR-*recB* $\Delta(TTAA...TGAT)_{36nt}$ | This work |

**Table S2:** Plasmids used in the study.

| Plasmids | Description | Source |
|---|---|---|
| pTOF24Δ*recB* | pTOF24-derivative plasmid used for construction of Δ*recB* strain | [32] [1] |
| pTOF24 *recB*-5′UTR | pTOF24-derivative plasmid used for construction of *recB*-5′UTR strain | This work |
| pBAD33 | Backbone plasmid used for construction of pIK02 | [33] |
| pIK02 | pBAD33-derivative plasmid carrying *recB* gene under control of arabinose-inducible promoter, P*araBAD* | This work |
| pZA21MCS | Backbone plasmid used for construction of pZA21-ChiX | Expressys |
| pZA21-ChiX | pZA21MCS-derivative plasmid carrying *chiX* | This work |
| pZA21-CyaR | pZA21MCS-derivative plasmid carrying *cyaR* | [27] |
| pDWS2 | pBR322-derivative plasmid carrying native *recC-ptrA-recB-recD* chromosomal region | [34] [2] |
| pQE80L | Backbone plasmid used for construction of pQE-Hfq | [35] [3] |
| pQE-Hfq | pQE80L-derivative plasmid carrying *hfq* | [35] [3] |
| pQE-Hfq(Y25D) | pQE80L-derivative plasmid carrying *hfq(Y25D)* | [35] [3] |
| pQE-Hfq(K56A) | pQE80L-derivative plasmid carrying *hfq(K56A)* | [35] [3] |

[1] Gift from Prof. David Leach, The University of Edinburgh
[2] Gift from Prof. Gerald Smith, The Fred Hutchinson Cancer Research Center
[3] Gift from Prof. Teppei Morita and Prof. Hiroji Aiba, Suzuka University of Medical Sciences

**Table S3:** Primers used for strain and plasmid construction.

| Primer ID | Sequence (5′-3′) |
|---|---|
| hfq_H1_P1 | GAATCGAAAGGTTCAAAGTACAAATAAGCATATAAGGAAAAGAG AGAATGGTGTAGGCTGGAGCTGCTTC |
| hfq_H2_P2 | CTCCCCGTGTAAAAAAACAGCCCGAAACCTTATTCGGTTTCTTCGC TGTCGGTCCATATGAATATCCTCCTTAG |
| ChiX_H1_P1 | TCTTGCCTAAGAGTATTGGCAGGATGGTGAGATTGAGCGACAATC GAGTTGTGTAGGCTGGAGCTGCTTC |
| ChiX_H2_P2 | CACCTGTATGGAGAAGGGAATTTGCCGCAAATGTTGCGCTAAAAA AATGGCGGTCCATATGAATATCCTCCTTAG |
| chiX_ZA21 | ACACCGTCGCTTAAAGTGACGGCATAATAATAAAAAAATGAAATT CCTCTTTGACGGGCCAATAGCGATATTGGCCATTTTTTTGGTACGCG TGCTAGAGGCATC |
| pZA21MCS_5P | 5P-GTGCTCAGTATCTCTATCACTGA |
| oIK03 | AAGCTTGGCTGTTTTGGCGGATGAG |
| oIK04 | ACCGAGCTCGAATTCGCTAGCCCAAAA |
| oIK10 | ACCCGTTTTTTTGGGCTAGCGAATTCGAGCTCGGTCCTGATGAGTG AAAAGAATGAGTG |
| oIK11 | GAAAATCTTCTCTCATCCGCCAAAACAGCCAAGCTTCTCCACAGCT TCCAGTAATTGC |

**Table S4:** The sequence of the gBlock used for the construction of the *recB*-5′UTR strain.

| |
|---|
| 5′-GTAATACAAGGGGTGTTATGAGCCATATTCAACGGGAAACGTCTTGCTCGAGGGTTCAG<br>CAATAGCATCGGCCAGGCGGTCTACCGCACCAGGCAAGGCGTCGTTCTCAACTTCCAGAT<br>AGAAAGCCGTGCGATACGGCGCAGTGCTGGCATTGTGACTACCGCCGTGCATTTTGAGAT<br>ATTCGGCCAGACTGTCAGCCTGCGGGTACTTTTTCGACCCCATCAGACTCATATGTTCAAG<br>GTAATGTGCCAGCCCCTGGTACGCCTCGGGATCTTCCAGCGACCCAACGGGCACCACCAG<br>CGCCGAGAGCGATTTAACTGCCTGCGGATCAGAAACCAGCAAGACCACCATACCGTTATC<br>CAGACGTATAGCCTGATACTGGCGGTTATCTTTATCACTTTTACGGATGGTTTCCTGAATCG<br>GCTGCCATCCCGTTTCTGCCTGACTTAAGGGTGCCCAAAGGGCAACTAACAACAATAATG<br>CTTTGAACCAGGTGCTGCGGGGCATTCACGGACCTCATAAGCTTCGCAAATCATCTGCCA<br>GAATTTAATCTTGTGCTGCACGAGTCAGCCTATGTTTATATAACCATCAGTCCGTGACTGGT<br>GCGCATCATAAAGTAAGCGGATAGATTGCGCAATTTTTATACAGCACTCATGACTGATTAA<br>AGCGAAACAGCGGTAACAGGAAACGTTGCGACTGTTCAACGATGGCCTCCATTGTCTCTG<br>GTGTTAATTGCCGCCAGAGCCTTTGATACCAGATATCATCACCTTCGCCACGCACCATCAT<br>GTTGCCTTCGTAAGCCTGAAGGAATTTCGTACGGGCTTTTTGCAACGTGGAATCGTCATCC<br>AGCATGGCATCGTTTTGCGCGTCATAACAGGTTTTTAGCCACGCGCCGCCACTTTCAGGTA<br>ACACCAGCAATGGCGCGGACATTCCTTCACGATACCCCTCAATCAGTTGTGAGAGGTAAT<br>GCAAAGCCTGTTCGGCGTCGACCGGTGGCGAATGGGACGCGCCCTGTAGCGGCGCATTA<br>AGCGCGGC-3′ |

**Table S5:** Oligos used for RT-qPCR quantification.

| **Sequence (5′-3′)** | **Gene** |
|---|---|
| TGGCCTGACGCGTATGTTGT | *recC* |
| TGCCCCACCAGTTCTGCAAT | *recC* |
| CTCTTGCGGGTTACGGACGT | *recD* |
| ATTCAGTCCAGCCACGCCAA | *recD* |
| TCTGGCTTCATCGCTCGCAA | *ptrA* |
| TGGGGCATGGGCTTCCTTTT | *ptrA* |
| ACCCGCGCATTGGCTGAGAT | *recB* |
| CACCGCTTTCGCTACGCAGC | *recB* |
| CGGTGGTCCCACCTGACC | *rrfD* |
| CCTACTCTCGCATGGGGAGACC | *rrfD* |
| CGTCTCGCCCGGTTTCTCAT | *hfq* |
| GGAAGTATTCTGCGCGCTGC | *hfq* |
| GTCGCTTAAAGTGACGGCAT | *chiX* |
| TCGCTATTGGCCCGTCAAAGA | *chiX* |
| GCTAGCTGTACCAGGAACCACC | *cyaR* |
| GGGAGATTACACAGGCTAAGGAGG | *cyaR* |

**Table S6:** Sequences of *recB* RNA FISH probes labelled with TAMRA dye.

| Probe ID | Sequence (5′-3′) |
|---|---|
| RecB-TAMRA_1 | ctgtaagggcaagcgcaaag |
| RecB-TAMRA_2 | gcaatcgtaaaggttttgcc |
| RecB-TAMRA_3 | tcgtggatattgctacggat |
| RecB-TAMRA_4 | gttcggctaacaacaaccac |
| RecB-TAMRA_5 | aaaggcattcaggttgagca |
| RecB-TAMRA_6 | aatcagctgctgctcaaaca |
| RecB-TAMRA_7 | aaagacgacctgggctattt |
| RecB-TAMRA_8 | cgccttgcagataacgatta |
| RecB-TAMRA_9 | ctgctgttttaccgtatcaa |
| RecB-TAMRA_10 | accagaagattcgatcagcg |
| RecB-TAMRA_11 | cggttaaacttgcgtcgatc |
| RecB-TAMRA_12 | atcttgtcgatccatttagc |
| RecB-TAMRA_13 | ccggcaactgataactgttt |
| RecB-TAMRA_14 | cttcgtgcgatcttctaaga |
| RecB-TAMRA_15 | ttgatcgatcgcctcaaaca |
| RecB-TAMRA_16 | cagatcgcggatcgacaatg |
| RecB-TAMRA_17 | agccgacttaacatgtcatc |
| RecB-TAMRA_18 | cgccaattagcaacaatgcg |
| RecB-TAMRA_19 | tacgcgccttcatataagtg |
| RecB-TAMRA_20 | tgtctaaagtgtagtgggcg |
| RecB-TAMRA_21 | aagcttattcacgctgttca |
| RecB-TAMRA_22 | tcgcgaaacatgaacgcgtc |
| RecB-TAMRA_23 | aaacggaagggatttccagc |
| RecB-TAMRA_24 | cctgcaacaaccaaagcatt |
| RecB-TAMRA_25 | cagatttgccgataaccatc |
| RecB-TAMRA_26 | gttttcagcaatgttacgcg |
| RecB-TAMRA_27 | ttgtagcagttcgctgatat |
| RecB-TAMRA_28 | tggatcgtgacaatctgcac |
| RecB-TAMRA_29 | tggacgcggaaattggtgat |
| RecB-TAMRA_30 | atcgtgataaaacgcctgct |
| RecB-TAMRA_31 | ttaagatccagaactgcctc |
| RecB-TAMRA_32 | aagcaaacgcagatcttccg |
| RecB-TAMRA_33 | aatgccaaaccgaacgtgtc |
| RecB-TAMRA_34 | aacgcttcaatacaggtgcg |
| RecB-TAMRA_35 | ttggttatcaccagtttgtg |
| RecB-TAMRA_36 | tcagctctgctgtagaaaca |
| RecB-TAMRA_37 | aaaccagagtagctggtgac |
| RecB-TAMRA_38 | tgatgtggtgttaacgtcgg |
| RecB-TAMRA_39 | tcaaccggctgggtaaaatc |
| RecB-TAMRA_40 | ttattgcgggcggaaagttg |
| RecB-TAMRA_41 | gataaaactccatctccacc |
| RecB-TAMRA_42 | aacgtatcaagctgactggc |
| RecB-TAMRA_43 | gccagggtataaagctgata |
| RecB-TAMRA_44 | caatgcgatggcgcagataa |
| RecB-TAMRA_45 | tggtgctcatagtcgtaatc |
| RecB-TAMRA_46 | acagataaataacgccgcca |
| RecB-TAMRA_47 | gatgttctttatcaacgcca |
| RecB-TAMRA_48 | cataccggcaaacatctcat |

**Table S7:** Parameters of the model of RecB expression.

| Parameter | Meaning | Value | 95% CI | Source |
|---|---|---|---|---|
| $k_m$ | transcription rate | 0.21 $min^{-1}$ | [0.13 0.33] $min^{-1}$ | Estimated |
| $\gamma_m$ | mRNA degradation rate | 0.62 $min^{-1}$ | [0.48 0.75] $min^{-1}$ | Measured |
| $b$ | mRNA burst size | 0.95 molec | [0.76 1.19] molec | Estimated |
| $k_p$ | translation rate | 0.15 $min^{-1}$ | [0.12 0.18] $min^{-1}$ | Estimated |
| $\gamma_p$ | protein removal rate | 0.015 $min^{-1}$ | [0.011 0.019] $min^{-1}$ | Measured |
